# Multiplexed high-throughput immune cell imaging reveals molecular health-associated phenotypes

**DOI:** 10.1101/2021.12.03.471105

**Authors:** Yannik Severin, Benjamin D. Hale, Julien Mena, David Goslings, Beat M. Frey, Berend Snijder

**Author notes:** Correspondence: Prof. Dr. Berend Snijder Institute of Molecular Systems Biology Department of Biology ETH Zürich Otto-Stern-Weg 3, building HPM office H44 CH-8093 Zürich, Switzerland, www.imsb.ethz.ch, www.snijderlab.org.

## Abstract

Phenotypic plasticity is essential to the immune system, yet the factors that shape it are not fully understood. Here, we comprehensively analyze immune cell phenotypes including morphology across human cohorts by single-round multiplexed immunofluorescence, automated microscopy, and deep learning. Using the uncertainty of convolutional neural networks to cluster the phenotypes of 8 distinct immune cell subsets, we find that the resulting maps are influenced by donor age, gender, and blood pressure, revealing distinct polarization and activation-associated phenotypes across immune cell classes. We further associate T-cell morphology to transcriptional state based on their joint donor variability, and validate an inflammation-associated polarized T-cell morphology, and an age-associated loss of mitochondria in CD4^+^ T-cells. Taken together, we show that immune cell phenotypes reflect both molecular and personal health information, opening new perspectives into the deep immune phenotyping of individual people in health and disease.

## Introduction

The morphology of a cell closely reflects its state, as it adapts to dynamic functional requirements and thereby constrains future behavior (Bakal et al. 2007; Folkman and Moscona 1978; Lecuit and Lenne 2007; Boutros, Heigwer, and Laufer 2015). This feedback mechanism has been shown to influence many cellular events, including cell differentiation (McBeath et al. 2004; Discher, Mooney, and Zandstra 2009), cell division (Carlton, Jones, and Eggert 2020; Ramkumar and Baum 2016; Folkman and Moscona 1978), adaptation to the microenvironment (Snijder et al. 2009; Snijder and Pelkmans 2011; Liberali, Snijder, and Pelkmans 2014), and malignant transformation (Hanahan and Weinberg 2011; Wu et al. 2020). Few differentiated healthy human cells change their phenotype as drastically as immune cells: a plasticity that is critical to the correct function of the immune system as a whole (Zhou, Chong, and Littman 2009; Galli, Borregaard, and Wynn 2011; Sica and Mantovani 2012). As a consequence, studying immune cellular heterogeneity at the molecular level has been transformative for our understanding of the immune system, measured for example by flow cytometry (Maecker, McCoy, and Nussenblatt 2012; Craig and Foon 2008), single-cell mass cytometry (Spitzer and Nolan 2016; Bendall et al. 2011), and single-cell RNA sequencing (Papalexi and Satija 2018; Jaitin et al. 2014; Shalek et al. 2013; Villani et al. 2017; Giladi and Amit 2018). Complementary to these molecular measurements, microscopy has shown the importance of immune cell morphology in multiple settings: distinct cellular morphologies are associated with, and influence the outcome of, monocyte polarization (Bertani et al. 2017; McWhorter et al. 2013) and T- and B-cell activation (Gómez-Moutón, Abad, and Mira 2001; K. B. L. Lin et al. 2008; van Panhuys, Klauschen, and Germain 2014; Russell 2008; Faure et al. 2004; W. Lin et al. 2015), and label-free imaging of hematopoietic cells has enabled predicting the outcome of future lineage choices (Buggenthin et al. 2017). Additionally, a recent study, using organelle marker abundance as a proxy for cell morphology, found extensive evidence for morphological heterogeneity in both healthy and diseased immune cells (Tsai et al. 2020). Due to their mixed adherent nature, however, primary immune cells such as peripheral blood mononuclear cells (PBMCs) were long considered incompatible with automated fluorescence microscopy, the tool of choice to characterize cellular morphology with spatial resolution across millions of cells (Snijder et al. 2009; Liberali, Snijder, and Pelkmans 2014; Perlman et al. 2004; Boutros, Heigwer, and Laufer 2015; Wawer et al. 2014; Young et al. 2008; Caicedo et al. 2017). This has hampered the comprehensive measurement and study of morphological heterogeneity present in the immune system, and thus has left unanswered the question of which molecular and health factors globally shape the compendium of human immune cell morphologies.

## Results

To be able to comprehensively measure immune cell phenotypes, we developed a multiplexed immunofluorescence approach for peripheral blood mononuclear cells (PBMCs) that extends our previously developed protocol for high-throughput image-based screening in human biopsies compatible with mixed non-adherent cells (Vladimer et al. 2017; Snijder et al. 2017; Kornauth et al. 2021) (Figure 1). In contrast to previously reported cyclical multiplexed immunofluorescence protocols (J.-R. Lin, Fallahi-Sichani, and Sorger 2015; Gerdes et al. 2013; Gut, Herrmann, and Pelkmans 2018), we stain once with a comprehensive immune cell marker panel that multiplexes 8 surface markers and a nuclear dye, which is imaged by automated confocal microscopy and brightfield imaging in a single run (Figure 1i and Supplementary Table 1). A deep convolutional neural network (LeCun, Bengio, and Hinton 2015) with custom architecture (Figure S1A) was subsequently used to classify each cell, making use of distinct marker expression patterns, lineage-specific labeling encoded by the staining panel, and likely differences in immune cell morphology (Figure 1i). The CNN was trained across eight immune cell classes, using 89’483 manually curated 5-channel sub-images (4 fluorescent channels and brightfield) centered on individual cells sampled from 15 healthy donors (available at https://doi.org/10.3929/ethz-b-000343106). The eight immune classes capture the predominant immune lineages present in PBMCs, including three distinct T-cell subsets (CD4^+^, CD8^+^ and CD4^-^CD8^-^), monocytes, dendritic cells, natural killer cells, B-cells, and nucleated immune cells negative for all 8 surface markers (Figure 1).

**Figure 1.**
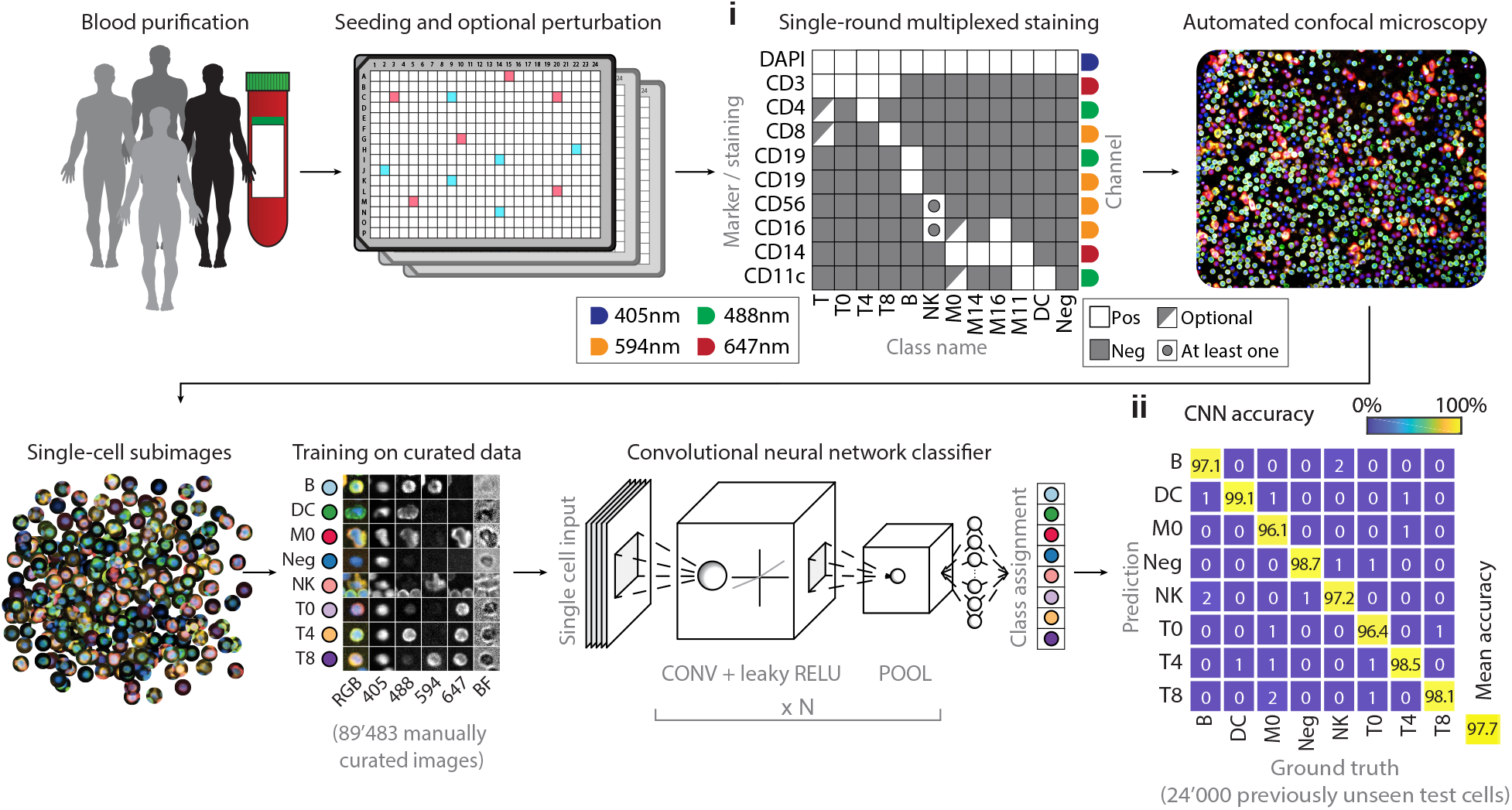
Workflow for the single-round multiplexed immunofluorescence, image-based screening, and associated deep learning-based classification of human peripheral blood mononuclear cells (PBMCs). PBMCs of healthy human donors are seeded in 384-well plates, optionally containing drugs or immune stimuli. Cells are fixed and stained with a comprehensive antibody panel (**1i**) and imaged by automated confocal microscopy. A convolutional neural network (CNN) is trained on 89483 manually curated sub-images to distinguish eight different immune cell classes, and subsequently classifies all cells in the experiment. The curated test set contains 100 cells per class per donor per staining condition. **1ii**, Confusion matrix of CNN performance across all 24’000 cells that the CNN did not see before.

CNN performance was stable across retraining, showed no sign of overfitting, and was 97% accurate for unseen donors systematically left out of the training data (Figure S1B and S1C). The network further achieved 97.7% classification accuracy (Figure 1ii) on a previously unseen test dataset of 24’000 curated cells comprising PBMCs from the same 15 healthy donors (Figure S1D). The classification efficiently demultiplexed the mixed marker signals such that the resulting abundances of each subpopulation matched our expectations (Figure S1E and F), and both the class fractions (Figure S1G) and class probabilities (Figure S1H) showed good reproducibility over different experimental replicates (median r = 0.90 and 0.95 respectively). Whilst marker expression likely contributed towards the classification accuracy between morphologically similar classes (such as T4 vs T8), cell morphology likely contributed to the separation of distinct cell types whose markers were multiplexed in the same channel, such as CD14+ monocytes and CD3+ T-cells both stained in the APC channel. Supporting this interpretation, a 2-class CNN could separate T-cells and monocytes with 95% accuracy based on just the label-free brightfield and DAPI channels (Figure S1I and J). Thus, the 8-class CNN learned to generalize immune phenotypes across individual donors and experiments, presenting a robust, efficient, and data-rich high-throughput screening strategy with broad applicability.

Both supervised and unsupervised deep learning algorithms are increasingly used for image clustering (Xie, Girshick, and Farhadi 2016; Aljalbout et al. 2018), which we here explored for the purpose of clustering immune cell phenotypes. The CNN returns a confidence vector for each cell that creates an 8-dimensional feature space, which we visualized by t-distributed stochastic neighbor embedding (t-SNE) (Maaten and Hinton 2008) (Figure 2A). To minimize possible batch effects and confounding factors from *ex vivo* culturing, we analyzed a subset of 10 of the 15 donors on which the CNN was trained, whose blood had been simultaneously processed, and incubated for just 1 hour before fixing and imaging across replicate wells and plates. Visualization of unperturbed immune cells from these 10 donors suggested considerable cell-to-cell variability, particularly among monocytes, even just within the cells classified with high CNN confidence (Figure 2A). Projecting molecular and morphological cell features measured by conventional image analysis on the t-SNE embedding revealed that the CNN had separated monocytes based on their CD16 and CD11c expression levels, even though it was not trained explicitly to do so (Figure 2A insert). Moreover, this showed that even for high-confidence cells the CNN class probabilities reflected marker expression and morphological heterogeneity for all 8 immune cell classes, with nuclear size and brightfield intensity differences observed within each class (Figure 2A and B). Thus, while the 8-class CNN was strictly trained in a supervised manner, its neural network uncertainty additionally allowed further grouping of previously unannotated cellular phenotypes, capturing recurrent phenotypes present in primary human immune cells.

**Figure 2.**
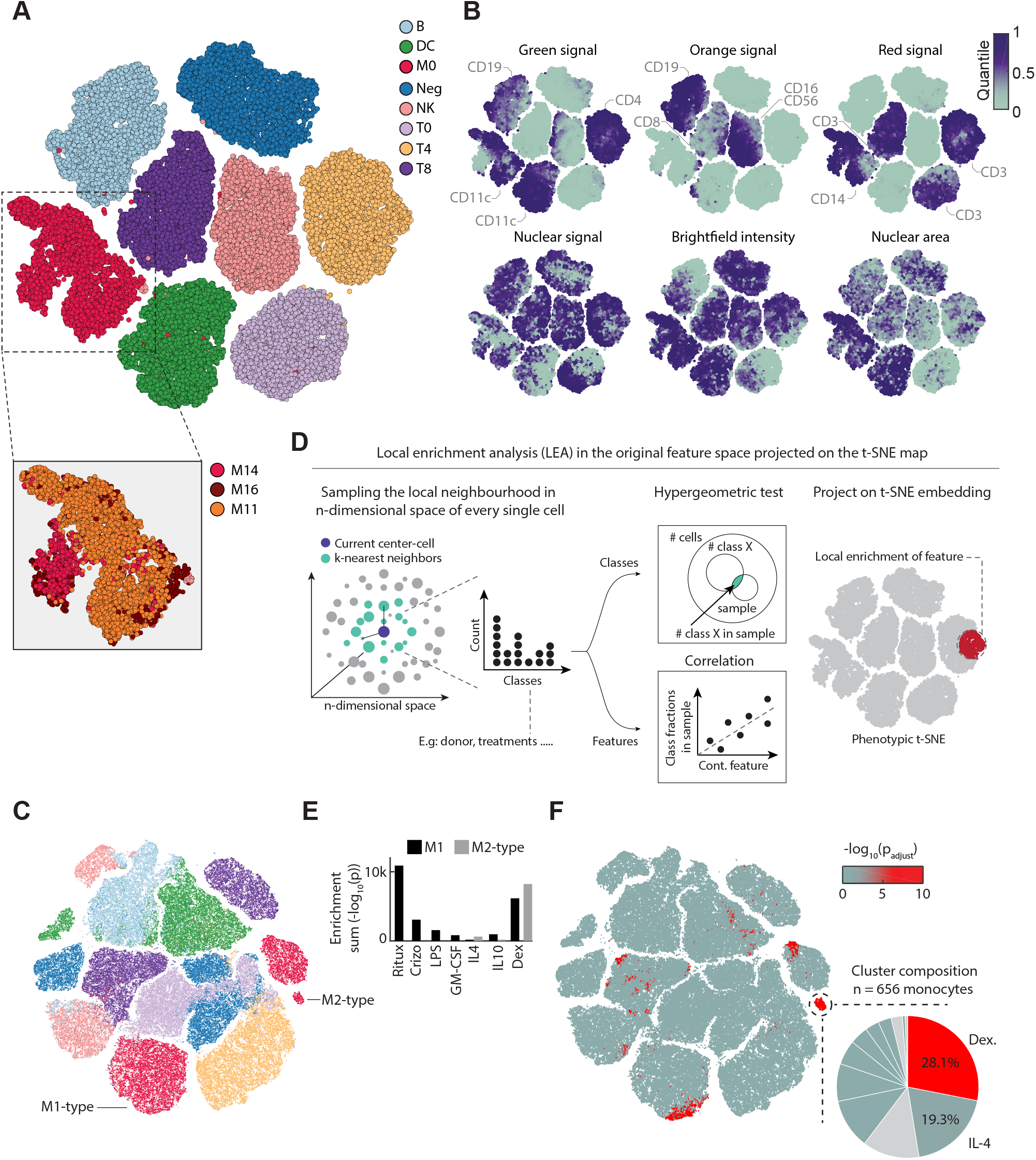
**(A)** Phenotypic landscape of the unperturbed immune system across ten healthy donors. t-Distributed Stochastic Neighbor Embedding (t-SNE) embedding of the 8-class CNN probabilities of up to 1000 randomly subsampled high-confidence multiplexed cells per class and per donor (class probability > 0.7). Monocytes are further divided into three subpopulations by thresholding the immunofluorescence (IF) intensity of CD16 and CD11c stainings, respectively (insert). Figure depicts a total of 78850 cells, randomly sampled from 40 wells for each of the 10 donors. All donors were processed and measured together in a single experiment, across 40 replicate wells per donor distributed over two 384-well plates. **(B)** Selected single-cell features projected onto the t-SNE shown in **2A**. Median value of overlapping data points is calculated and color is assigned accordingly. Points are plotted in order of intensity, with the lowest intensity on top. **(C)** Phenotypic landscape of the *ex vivo* perturbed immune system of a single donor. The CNN class probability t-SNE map on the left shows 600 randomly chosen single cells per cell class and drug treatment, colored by class assignment. **(D)** Overview of the local enrichment analysis (LEA) workflow. LEA probes the k-nearest neighbors of each single cell in a multidimensional space for enrichment of either continuous or discrete features. For discrete features, the baseline probability of finding n cells of condition X in the probed neighborhood follows a hypergeometric distribution, from which an enrichment p-value is calculated (taking into account the total number of drawn cells, the total number of cells in the t-SNE and the total number of cells of condition X in the dataset). For continuous features, the relative fraction of cells of each donor in the probed local neighborhood is calculated. These fractions are then rank-correlated with a continuous feature that was measured across donors. The enrichment probability for continuous features corresponds to the p-value of the correlation. In both cases, the enrichment probability is assigned to the center-cell and the approach is iterated for each single cell in the analysis. **(E)** Bar graph depicting the sum total log10(LEA P-values) for selected perturbations in the M1-type (black bars) and M2-type (grey bars) monocyte clusters. **(F)** LEA analysis reveals regions in the phenotypic space that are significantly enriched for dexamethasone-treated PBMCs. Cells in the t-SNE embedding are colored by their enrichment significance of the LEA analysis run in the original 8-class probability space (−log_10_(p_adjust_); see colorbar). Insert highlights the contribution of different perturbations to the selected M2-type monocyte cluster. Figure depicts a total of 199375 cells, randomly sampled from across 240 wells for a single donor.

We next tested if this deep learning uncertainty could also be used to quantify and categorize extrinsically induced changes in immune cell phenotypes. To this end, we stimulated PBMCs from a single donor with 12 immune modulators *ex vivo* across concentrations and replicates, measuring 5 million multiplexed stained and imaged PBMCs (Supplementary Table 2). First, we visualized the structure in the CNNs confidence by t-SNE (Figure 2C), equally sampling cells from across all 8 classes and 12 perturbations. This revealed monocytes to be divided into three clusters associated with distinct CNN confidence profiles, not trivially explained by marker expression differences (Figure S2A, B and C). To identify the contribution of distinct immune modulators to the morphological landscape of immune cells, we developed a method called K-nearest neighbor local enrichment analysis by hypergeometric testing (LEA, Figure 2D and methods). For each cell, LEA identifies the nearest neighbors in the original 8-class probability space and calculates the hypergeometric significance of enrichment for cells with a certain property in this neighborhood. LEA next assigns this significance back to the original starting cell. Projecting the LEA results back on the t-SNE embedding revealed that the monocyte subcluster with the lowest CNN confidence were enriched for monocytes exposed to M1-type inducing agents *E. coli* lipopolysaccharides (LPS) and GM-CSF (Figure 2E and Figure S2D) (Martinez and Gordon 2014), or cytotoxic agents causing the release of danger-associated molecular patterns. The second monocyte cluster was strongly enriched for cells exposed to M2-type associated dexamethasone or IL4, while the third, highest confidence, monocyte cluster was not enriched for most perturbations, thus likely reflecting unperturbed monocyte phenotypes (Figure 2F).

Stimulation with microbial compounds like LPS can selectively alter immune cell crosstalk, for example through the induction of cell-cell contacts. We therefore suspected that phenotypes in the M1-type cluster could in part reflect changes in the multi-cellular context. To verify this, we performed spatially resolved single-cell analysis across the 8 classified immune cell types, allowing the high-throughput screening of 36 distinct immune cell-cell interactions simultaneously, a significant increase compared to our previous non-multiplexed efforts (Vladimer et al. 2017) (Figure S3A-B). Indeed, analysis of all 43 million cell-cell interactions measured in this experiment (Figure S3A) confirmed the M1-like monocyte cluster to be enriched for monocyte-to-monocyte interactions (Figure S3C). Thus, LPS-mediated monocyte activation led to distinct M1-like monocyte phenotypes, defined in part by an altered multi-cellular context. Collectively, LEA revealed that the uncertainty of the deep neural network reconstituted previously established monocyte M1/M2-type polarization phenotypes in a fully unsupervised manner (Fig 2), while exposing considerably phenotypic complexity, with most immunomodulatory perturbations simultaneously affecting the phenotype of multiple immune cell class (Figure 2C and Figure S2D).

The phenotypic heterogeneity of circulating immune cells captured by our image-based measurements could reflect both genetic and non-genetic influences (Melé et al. 2015; Galli, Borregaard, and Wynn 2011). To explore this we analyzed commonalities and differences in the unperturbed immune phenotypes across the discovery cohort of the 10 donors shown in Figure 2. We first used LEA to measure enrichment of cells from the same donor in the nearest-neighborhood in the 8-dimensional CNN class probability space. This identified distinct cellular phenotype-regions significantly enriched for each of the 10 donors across several immune cell classes (Figure 3A). As these enriched phenotypes were measured across technical repeats, they potentially indicated donor-individual characteristics of immune cell morphologies, but could also reflect batch effects acting upstream of our sample processing and imaging. Repeating the analysis with randomized donor labels and comparing the sum of enrichments showed that the actual donor-enrichment in nearest neighbors of the latent space was well above what would be expected by random (P < 1.1x10^-308^; Figure 3A insert). We next looked for phenotypes that were enriched in donors with the same biological gender, with the 10 donors including 4 women and 6 men. This revealed strong gender associations with various immune cell morphologies (P < 1.1x10^-308^; Figure 3B), with NK- and Negative-cell class phenotypes particularly enriched in female donors, and not explained by enrichment in any individual female donor (Figure S4A).

**Figure 3.**
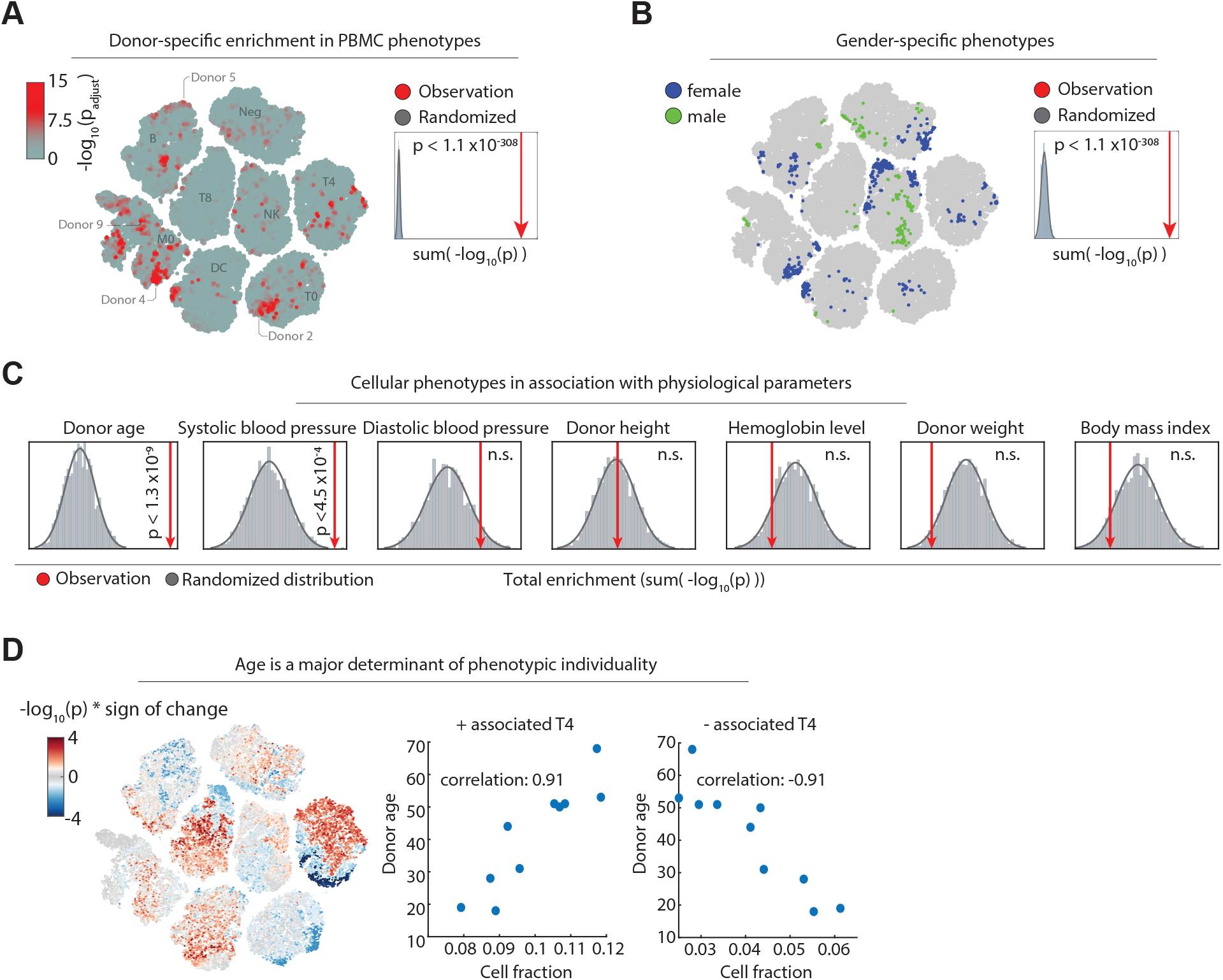
**(A)** LEA of donor-specific cells across 10 donors, visualized on the t-SNE of Figure 2A. The cells are colored by their maximum LEA significance across the 10 donors (−log_10_(p_adjust_); see colorbar). Insert: A randomized null distribution of donor enrichments was generated by randomizing the donor labels 2000 times and summing up all single-cell enrichments calculated by LEA per randomized run (grey bars). Sum enrichment of the actual data is shown in red, and the significance compared to the randomized runs is calculated by a one-sided t-test. **(B)** LEA of biological gender-specific phenotypes projected onto the t-SNE embedding. Cells are colored by their significant enrichment in female (blue) or male (green) specific phenotypes. Insert: A null distribution of random gender enrichment (grey) was generated by randomizing the donor labels 2000 times and summing up all single-cell enrichments calculated by LEA. Sum enrichment of the actual data is shown in red, as in 3A. **(C)** Association analysis of various health parameters with cellular phenotypes calculated by LEA. Null distributions of random correlation significance (grey) were generated by randomizing the donor labels 2000 times and summing up the all single-cell enrichments calculated by LEA per randomized run. Enrichment of the actual data is shown in red (one-sided t-test). **(D)** LEA Age-associations projected onto the t-SNE embedding. Single-cells are colored by their signed significance of correlation (−log_10_(p) * sign of the correlation; see colorbar). Insert: Fraction of all significantly positive and negative age-associated CD4^+^ T-cells with donor age (p<0.05).

We next explored immune phenotype associations with continuous health parameters such as donor age, which has been described to dramatically alter the immune phenotypic landscape (Carr et al. 2016; Alpert et al. 2019) (Figure 3C, Figure 2D and methods). A modification of LEA for continuous variables calculates the significance of the rank correlation between the fraction of cells per donor in the nearest neighborhood and any continuous variable of each donor (Figure 2D). As before, the LEA analysis was run in the 8-dimensional CNN class probability space. To correct for spurious associations, we compare the association strength with those observed in many repeats with the same health parameter randomized across the donors. Testing donor age, height, weight, body mass index, blood pressure, and hemoglobin levels revealed significant associations with donor age (P < 1.3 x 10^-9^) and systolic blood pressure (P < 4.5 x 10^-4^; Figure 3C), but not to any of the other measured health parameters. The age-associated phenotype map revealed bimodal age associations for several immune subpopulations, particularly striking for CD4^+^ T-cells (Figure 3D). Across the cells that make up the phenotype map, the age associations were mutually exclusive of the single donor enrichments (r = -0.002; Figure S4B).

To investigate the above identified phenotypic and health associations we next used LEA to associate molecular pathway expression as measured by transcriptomics with immune cell phenotypes. Focusing on T-cells, we performed bulk RNA-sequencing of CD3 positive cells purified from the same 10 healthy donor blood samples, detecting on average around 15’000 expressed transcripts (Figure S5A). LEA rank-correlated local phenotype abundance (in the 8-dimensional CNN class probability space) with transcript abundance, analyzing T-cells randomly subsampled from each donor to match the population composition measured by RNA-sequencing (Figure 4A). To benchmark these phenotype-to-transcriptome associations, we first compared the LEA associations of *CD4* and *CD8A* transcript abundance (Figure 4A) with the CD4 and CD8 protein expression levels explicitly measured by immunofluorescence for each T-cell (Figure S5B). Validating the approach, LEA achieved excellent results for these proof-of-concept benchmarks, with areas under the receiver operating curve of 0.93 and 0.89 for CD4 and CD8 positive cells, respectively (Figure 4B).

**Figure 4.**
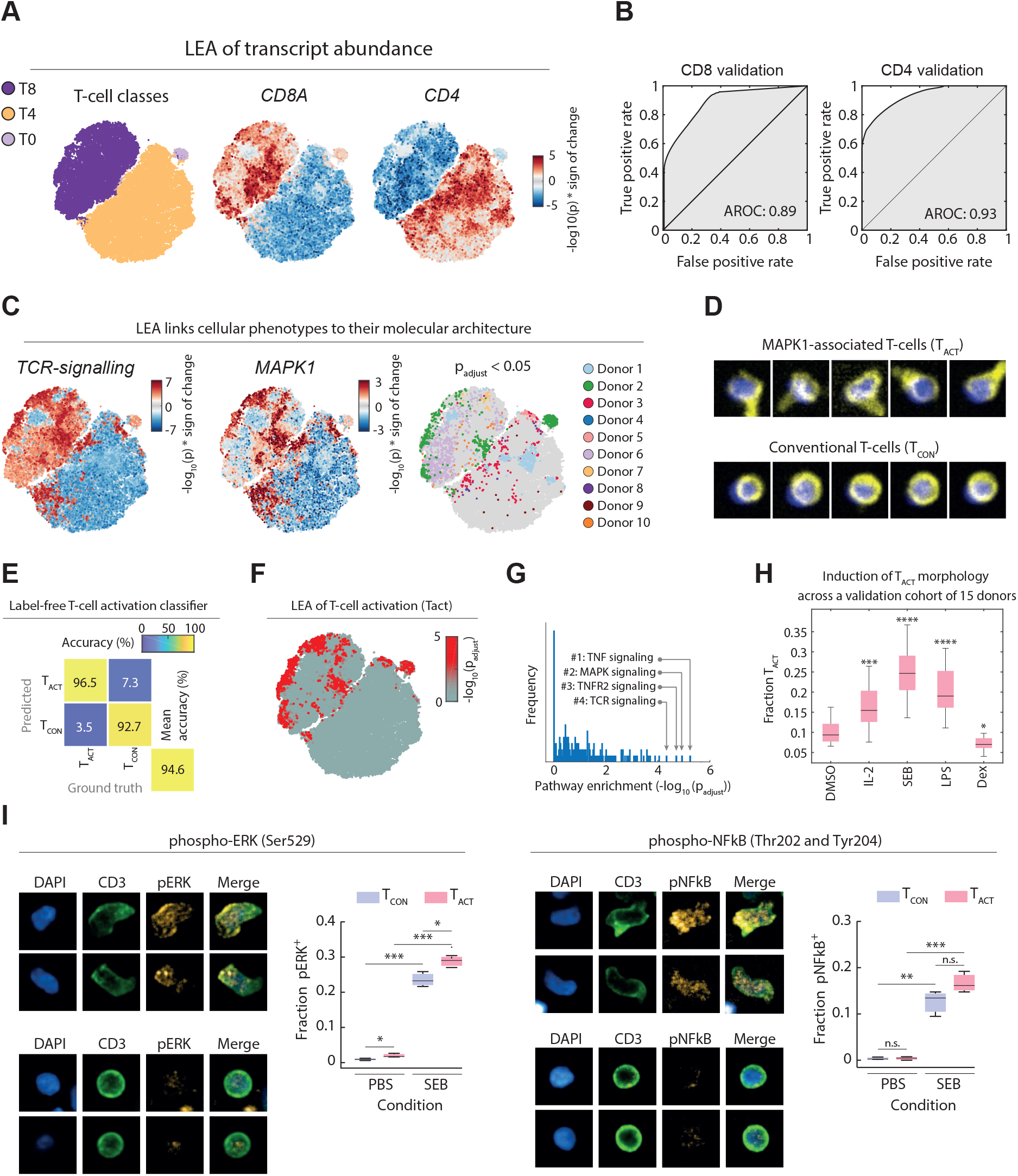
**(A)** t-SNE embedding of the CNN class probability across T-cells of ten healthy donors. 10000 T-cells are shown per donor, selected without confidence threshold to reflect the original abundance of T0, T4, and T8 subpopulations per donor. Left panel: t-SNE color-coded per T-cell class (see legend); Middle panel: t-SNE map colored by LEA-based associations with *CD8A* transcript abundance; Right panel: t-SNE map colored by LEA-based associations with *CD4* transcript abundance (−log_10_(p) * sign of the correlation; see legend). **(B)** Receiver operating characteristic (ROC) curves for the consistency between the *CD8A* transcript abundance LEA associations with CD8 expression levels by IF (left panel), and the *CD4* transcript abundance LEA associations with CD4 expression levels by IF (right panel). **(C)** LEA-based associations of TCR-signalling (left), *MAPK1* transcript abundance (middle) and donor enriched regions (right, p_adjust_ < 0.05 colored per donor) projected onto the T-cell phenotype map of **A. (D)** Examples of morphologically representative CD8^+^ T-cells from the positively MAPK1-associated regions (T_ACT_), and conventional CD8^+^ T-cells of other regions (T_CON_). Crops are 15 x 15 µm in size. Yellow = CD3, blue = DAPI. **(E)** Confusion matrix assessing the accuracy of the label-free T-cell activation (T_ACT_) classifier. The test set comprises 369 randomly selected T_ACT_cells, and 738 randomly selected T_CON_cells across multiple donors (including the 10 depicted donors). **(F)** LEA of the T_ACT_ phenotype projected on the t-SNE map (see colorbar). **(G)** Distribution of pathway significance across all retro-actively classified T_ACT_ cell morphologies. Pathway enrichments were calculated using a hypergeometric test on positively associated genes (top 0.95 percentile), and p-values were corrected for multiple testing. Significance of the top four most enriched pathways are indicated by grey arrows. **(H)** Induction and suppression of the T_ACT_ phenotype with immunomodulatory agents across an independent validation cohort of 15 individual donors. All compounds were screened at a concentration of 100ng/ml. Boxplots show the mean relative fraction of T_ACT_ cells in the T-cell compartment across all wells of each condition per donor. Stars indicate significance of T_ACT_ fraction per condition, compared with controls calculated with an unpaired t-test. **(I)** Immunofluorescence quantification of phospho-NFkB and phospho-ERK levels in T_ACT_ and T_CON_ cells. Boxplots show the fraction of phospho-signaling marker positive T_ACT_(red) and T_CON_cells (blue) after 48h incubation in the presence or absence of SEB. Boxplots show distributions of three technical repeats. Images show representative T_ACT_ and T_CON_ cell morphologies at 40X magnification. Crops are 15 x 15 µm in size.

We next sought to validate these pathway-phenotype associations by querying the associations the other way round: Starting from well-known pathways, and seeing what phenotypes are associated with it. To this end we inspected the associations with the T-cell receptor (TCR) signaling pathway as proxy for T-cell activation. TCR-signaling was strongly associated with distinct subregions of the phenotype map, including the cluster-periphery of CD8^+^ T-cells (Figure 4C). This pattern was recapitulated by the LEA associations with *MAPK1* (*ERK2*), part of the TCR-induced signaling cascade, which largely, but not exclusively, overlapped with regions enriched for cells from Donor 2 (Figure 4C). Visual inspection of cells residing in TCR-signaling and *MAPK1-*associated phenotypic regions revealed a striking polarized and activated T-cell morphology, henceforth referred to as T_ACT_ cells. In contrast, randomly sampled cells from adjacent and non-enriched regions contained conventional small and round T-cell morphologies, which we refer to as T_CON_ cells (Figure 4D). To robustly quantify the T_ACT_ morphology further, we trained a dedicated CNN on manually curated T_ACT_ and T_CON_ phenotypes, which achieved 94.6% validation accuracy on images from donors and experiments it was not trained on (Figure 4E and S5C). This allowed us to retroactively detect the T_ACT_ morphology for all imaged T-cells, which confirmed that the phenotype was present in all donors, and most enriched in the cells of Donor 2 (Figure 4F and Supplementary Figure S5D). Coming full circle, the T_ACT_ enriched regions associated with tumor necrosis factor (TNF) and MAPK-signaling as most-enriched pathways after multiple testing correction (Figure 4G).

To confirm that the T_ACT_ morphology is associated with inflammation and T-cell activation in an independent validation cohort, we stimulated PBMCs derived from 15 additional healthy donors with pro-inflammatory cytokine IL-2, superantigen Staphylococcus aureus Enterotoxin B (SEB), or LPS, which all led to significant increases in the fraction of T-cells adopting a T_ACT_ morphology (Figure 4H and S5E). Exposure to the anti-inflammatory synthetic glucocorticoid Dexamethasone, in contrast, reduced the relative abundance of T_ACT_ cells across the 15 donors (Figure 4H and S5E). To rule out the possibility that the T_ACT_ morphology was induced by cellular fixation prior to imaging, we further conducted live cell imaging of SEB stimulated PBMCs and visually confirmed the induction of the T_ACT_ cell phenotype (Figure S5F). We next measured by immunofluorescence the levels of phosphorylated NFkB (Ser529) and ERK (Thr202 and Tyr204) as a function of T-cell morphology, at baseline and upon SEB-stimulation in PBMCs. At baseline, T_ACT_ cells showed slightly but significantly higher levels of phosphorylated ERK. SEB-stimulation increased phosphorylated levels of ERK significantly higher in T_ACT_ than T_CON_ cells. Taken together, these results experimentally validated the LEA-based pathway enrichment analysis with the polarized T_ACT_ morphology. Thus, part of the donor unique fingerprints we previously observed had resulted from differences in T-cell activation between the donors, with 15% of T-cells from Donor 2 adopting the T_ACT_ morphology, predominantly in CD8^+^ T-cell compartment, while on the other end of the spectrum, only 7% of Donor 1 T-cells were T_ACT_ cells, here mostly in CD4^+^ T-cells (Figure S5D).

Having validated the phenotype-to-pathway association approach and its ability to discover and correctly describe new cellular phenotypes, we explored the pathway enrichments for age-associated T-cell phenotypes (Figure 5A and S6A). Pathways enriched in phenotypes that were reduced with age included nucleotide excision repair, telomere maintenance (Roth et al. 2003), cilia assembly (Stephen et al. 2018) and propanoate metabolism (Figure S6A). In contrast, pathways associated with T-cell phenotypes that increased with age included inflammation and stress-related pathways, particularly for the CD8^+^ compartment, and lysosome and vesicle-associated pathways in CD4^+^ T-cells (Figure 5A right). Inflammation is a well described risk factor for age-associated diseases (Franceschi, Bonafè, and Valensin 2000), and, consistently, the age-associated phenotypes overlapped partially with the above validated phenotype for activated CD8^+^ T-cells (Figure 5A right). Furthermore, impaired organelle and lysosome homeostasis in aged CD4^+^ T-cells has been previously described as a relevant process in aging of T-cells (Jin et al. 2020).

**Figure 5.**
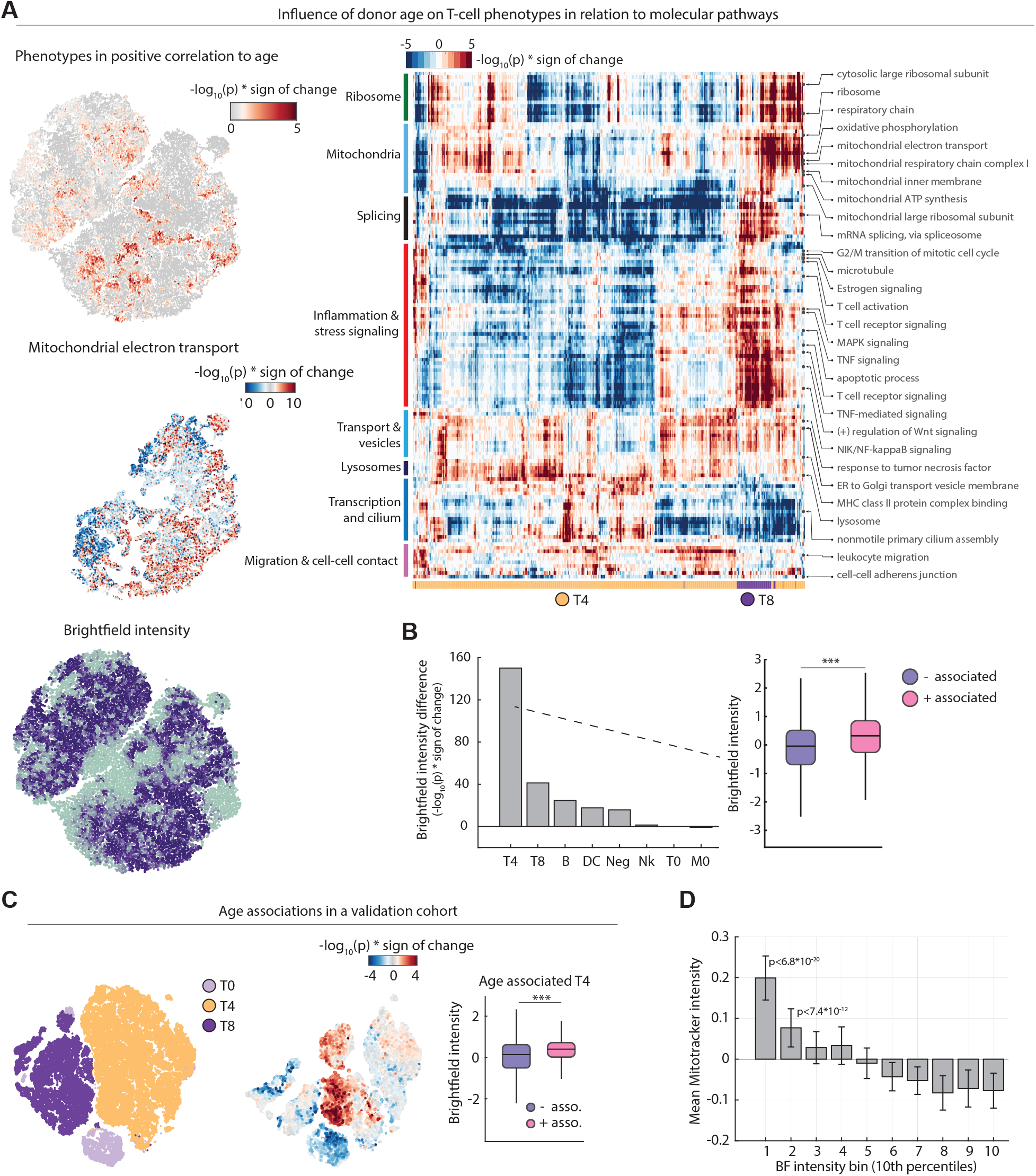
**(A)** Upper left: positive LEA associations with donor age projected on the t-SNE embedding as in Figure 4A (colored by -log_10_(p)). Middle left: LEA associations of Mitochondrial electron transport projected on the t-SNE embedding. Lower left: Brightfield single-cell intensity projected onto the t-SNE embedding. Median value of overlapping data points is calculated and color is assigned accordingly. Points are plotted in order of intensity, with the lowest intensity on top. Right: Heatmap overview of all significantly enriched pathways in positive age-associated T-cells (-log_10_(p)>5). **(B)** Comparison of the significance in difference in brightfield intensity of positively- vs negatively-associated immune cells per population (with an association cutoff of -log_10_(p)>1.3). Bar plots show the -log_10_(p) * sign of the change (1-(median(positive enrichment) / median (negative enrichment))). **(C)** Negative and positive age associations with cellular T-cell phenotypes and donor age in an independent validation cohort of 15 healthy individuals calculated by LEA (left and middle). t-SNE depicts a total of 5000 cells per donor. Right: Comparison of differences in brightfield intensity of positively vs negatively age-associated CD4^+^ T-cells (with an association cutoff of -log_10_(p)>1.3). **(D)** Mitochondrial content (as measured by MitoTracker) of CD4^+^ T-cells decreases with increased brightfield intensity. Bar-plots display the mean MitoTracker intensity of CD4-T-cells per well per 10-percentile bins of BF intensity within each well. Mean and standard deviations across 10 repeat wells with a combined total of n=78095 CD4+ T-cells are shown. P-values are from a two-tailed t-test of all replicate wells per bin against those of the brightest BF(right most) bin.

Pathway enrichments for oxidative phosphorylation and mitochondrial respiration in age-associated T-cell phenotypes were in line with reports of defective respiration in CD4^+^ T-cells of aged mice (Ron-Harel et al. 2018; Gomes et al. 2013), and suggested that the neural network might have identified a phenotypic T-cell signature associated with both donor age and mitochondrial abundance. Interestingly, the CD4^+^ T-cells showed strong brightfield intensity differences, a measure of intracellular granularity (Figure 2A,B and S5B). This brightfield-trend followed the age-associations we observed, with CD4^+^ T-cells enriched in younger people measured to be more granular (referred to as T4_BFD_ for ‘brightfield dark’ CD4^+^ T-cells; Figure 2A,B and 3C). Quantifying this association across all subpopulations, CD4^+^ T-cells indeed showed the most significant age-associated brightfield intensity differences (P < 10^-70^), followed by the CD8^+^ T-cells (P < 10^-40^), and less for the other immune cell classes (Figure 5B).

To reproduce this association we sampled an additional validation cohort of 15 healthy donors (Figure 5C), and trained a different neural network architecture on a new set of images generated only from this validation cohort (Figure S6B). This independent repetition of the workflow revealed that the age-associated T4_BFD_ phenotype was independent of the donor cohort and neural network and experimental batch (Figure 5C and Figure S6B). The age-associated brightfield intensity differences and mitochondrial pathway association might reflect loss of mitochondrial abundance in age in CD4^+^ T-cells (Murera et al. 2018). To support this interpretation we analysed if BF intensity reflects mitochondrial abundance using the natural heterogeneity observed within CD4^+^ T-cells of a single donor (Figure 5D). Indeed, those cells that were darkest by brightfield imaging displayed significantly higher mitochondrial abundance as measured by image-based quantification of the MitoTracker dye (Figure 5D). The deep learning uncertainty thus had revealed a label-free phenotype reflecting an age-associated mitochondrial decline in CD4^+^ T-cells, explaining in part how immune cell phenotypes measured by our high-throughput single-cell imaging pipeline capture donor information such as age.

## Discussion

We here explore the molecular health determinants of human immune cell phenotypes using a workflow that combines automated high-throughput microscopy, single-round multiplexed immunofluorescence, and deep learning-based phenotypic analysis. The presented method for phenotyping of immune cells distinguishes itself for its ability to integrate cell morphology, protein levels and localization, and multi-cellular context into a quantitative metric across 8 major immune cell classes, hundreds of conditions, and millions of cells. The resulting single-cell phenotype space, derived from the CNN’s uncertainty, reflected both genetic and non-genetic donor health information. We find age, gender, blood pressure, and inflammatory state to be significantly associated with human immune cell phenotypes, yet many more influences likely exist and more phenotype-associations captured by our approach remain unexplored.

Our workflow is tailored to make use of two large sources of biological heterogeneity: the heterogeneity observed between individuals, and heterogeneity observed within cells of the same class and donor. That dependency however is at the same time its limitation: The single-round multiplexed staining strategy benefits from the presence of multiple cell types with variable cell morphologies and marker profiles, and LEA requires donor or condition heterogeneity to power its associations. Furthermore, while the marker panel shown here reliably captures the predominant immune cell classes present in PBMCs, it does not resolve certain smaller subpopulations, such as Natural Killer T-cells (Bendelac, Savage, and Teyton 2007). However, the approach is flexible as the panel composition can readily be tailored to the identification of additional subpopulations, or adapted to different tissues, building on the same logic developed here.

Whilst this is not the first work which deploys CNN-based cell classification (Moen et al. 2019; Kraus et al. 2017; Kraus, Ba, and Frey 2016; Pärnamaa and Parts 2017; Dürr and Sick 2016; Sommer et al. 2017; Kandaswamy et al. 2016; Godinez et al. 2017; Hussain et al. 2019) and feature extraction (Pärnamaa and Parts 2017; Kraus et al. 2017; Jackson et al. 2019; Godinez et al. 2017), to our knowledge, this is the first work where deep learning is applied in high-throughput screening and phenotypic analyses of primary human PBMCs. By training the CNN on curated cells from across independent experiments, multiple donors, and conventional and multiplexed staining panels, we could prevent overfitting on phenotypes of single donors and technical bias stemming from experimental conditions. However, the CNN class probability space, which we here successfully employ as a phenotype discovery tool, is sensitive to different phenotypes resulting from different experimental conditions. As such, while CNN classification can be trained to be robust, experimental care needs to be taken when interpreting the CNN class probability space.

Once new phenotypes are discovered, as we demonstrate for the inflammation-associated T_ACT_ cell morphology, the ability to retroactively re-classify cells based on their morphology with dedicated CNNs allows robust morphological sub-classification of previously imaged cells even in absence of tailored marker panels. Attesting to the robustness of the discovered phenotypes, the inflammation-associated T_ACT_ and age-associated T4_BFD_ phenotypes could be validated in independent experiments, in an independent validation cohort, using distinct neural network architectures, and, for the T_ACT_ morphology, in both live-cell and fixed sample imaging.

In the future, repeated profiling of individual donors will allow to further stratify temporally stable from dynamic immune cell phenotypes. Furthermore, comparative studies across larger patient and donor cohorts, and identifying clinically relevant cell morphologies in the context of personalized treatment identification for hematological malignancies (Snijder et al. 2017; Kornauth et al. 2021), will be additionally attractive avenues of study. This will inevitably define the boundaries of the personal health information reflected by immune cell phenotypes. Given that the workflow allows simultaneous phenotype discovery combined with the molecular and personal health associations, it is well positioned to lead to the discovery of more as yet undescribed and clinically relevant immune cell phenotypes.

## Methods

### Experimental model

Buffy coats or whole blood tubes were obtained from coded healthy donors provided by the Blutspende Zurich, under a study protocol approved by the cantonal ethical committee Zurich (KEK Zurich, BASEC-Nr 2019-01579). Detailed donor information can be found in Supplementary Table 3.

### Experimental details

#### Collection and purification of human peripheral blood mononuclear cells (PBMCs)

Buffy coats or whole blood tubes were obtained from coded healthy donors provided by the Blutspende Zurich, under a study protocol approved by the cantonal ethical committee Zurich (KEK Zurich, BASEC-Nr 2019-01579). Healthy donor buffy coats or blood samples were diluted 1:1 in PBS (Gibco) and PBMCs were isolated with a Histopaque-1077 density gradient (Sigma-Aldrich) according to the manufacturer’s instructions. PBMCs at the interface were collected, washed once in PBS and resuspended in media. In all experiments, immune cells were cultured in RPMI 1640 + GlutaMax medium (Gibco) supplemented with 10% fetal bovine serum (FBS, Gibco) and incubated at 37°C with 5% CO_2_. Cell number and viability was determined utilizing a Countess II Cell Counter from Thermo Fisher according to the manufacturer’s instructions.

#### Non-adherent PBMC monolayer formation and drug screening and cell fixation

In the proof-of-concept drug screen, 5μl of a selected screening compounds (10x stock), and all respective controls (as outlined in Supplementary Table 2) were transferred to CellCarrier 384 Ultra, clear-bottom, tissue-culture-treated plates (PerkinElmer) with five replicates per condition. All conditions were screened in four concentrations: Cytokines (0.1, 1, 10, 100ng/ml); Rituximab (0.05, 0.1, 0.5, 1μg/ml); LPS (0.1, 1, 10, 100 ng/ml); Dexamethasone (0.4, 4, 40, 400ng/ml); Crizotinib (0.01, 0.1, 1, 10μM). 50 μl of medium containing approximately 4*10^5^ cells/ml was pipetted into each well of a 384-well compound plate and cells were allowed to settle to the bottom. The whole blood samples of the discovery cohort (shown in Figure 2A-B, Figure 3-5) were incubated for 1h, whereas all buffy coat samples, including all samples from the validation cohort (Figure 4H and Figure 5C) were incubated for 24 hours. All assays were terminated by fixing and permeabilizing the cells with 20μl of a solution containing 0.5% (w/v) formaldehyde (Sigma-Aldrich), 0.05% (v/v) Triton X-100 (Sigma-Aldrich), 10mM Sodium(meta)periodate (Sigma-Aldrich) and 75mM L-Lysine monohydrochloride (Sigma-Aldrich), for 20 minutes at room temperature. For Mitotracker staining (Thermo Fisher), cells were stained live with 500nM Mitotracker Red, prior to fixation. Fixative-containing medium was subsequently removed, and cells were blocked and photobleached in 5% FBS/PBS overnight at 4°C. Photobleaching was used to reduce background fluorescence and was performed by illuminating the fixed cells with conventional white light LED panels.

#### Immunostaining and Imaging

All fluorescent primary antibodies utilized in this work (outlined in Supplementary Table 1) were used at a 1:300 dilution in PBS. All antibody cocktails for immunohistochemistry (IHC) contained 6µM DAPI (Sigma-Aldrich) for nuclear detection. Before IHC staining, the blocking solution was removed and 20μl of the antibody cocktail was added per well and incubated for 1h at room temperature. Besides fully-multiplexed wells, each plate additionally contained several staining-control wells with a reduced number of antibodies (Supplementary Table 1). The staining-control wells served for evaluating antibody functionality and the generation of the CNN-training data (see below). For imaging, a PerkinElmer Opera Phenix automated spinning-disk confocal microscope was used. Each well of a 384-well plate was imaged at 20× magnification with 5×5 non-overlapping images, covering the whole well surface. The images were taken sequentially from the brightfield (650-760 nm), DAPI/Nuclear signal (435-480 nm), GFP/Green signal (500-550 nm), PE/Orange signal (570-630 nm) and APC/Red signal (650-760 nm) channels. Subsequently, the raw .tiff images were transferred from the microscope for further analysis.

#### Conventional image analysis and quality filtering

Cell detection and single-cell image analysis was performed using CellProfiler v2 (Carpenter et al. 2006). Nuclear segmentation was performed via thresholding on DAPI intensity. Cellular outlines were estimated by a circular expansion from the outlines of the nucleus. Additionally, a second and larger expansion from the nuclei was performed to measure the local area around each single cell (local cellular background). Standard CellProfiler based intensity-, shape- and texture features of the nucleus, cytoplasm and the local cell proximity were extracted for each measured channel. Raw fluorescent intensities were log_10_ transformed and normalized towards the local cellular background as described in Vladimer et. al., 2017 (Vladimer et al. 2017).

#### Convolutional Neural Networks

Convolutional neural networks used in this work were implemented using *MATLAB’s Neural Network Toolbox Version R2020a*. The curated dataset used in training, validation and testing of the CNN framework contains images of cells from fully multiplexed stainings and images from staining controls. Staining controls were designed to contain only a subset of the antibodies used in the multiplexed setting (Supplementary Table 1). This reduced complexity first enables to evaluate the functionality of the selected antibody and the presence of the targeted antigen in each sample. Furthermore, antibody combinations in the staining-controls were picked to mirror the staining of the selected subpopulation in the multiplexed setting (e.g. staining-control 1 only contained antibodies marking T-cell specific antigens; T-cells in the multiplexed setting will have the same staining pattern). The same staining patterns in the controls and the mostly-non-overlapping emission spectra of the chosen antibodies allow an easy, marker-intensity-based identification of subpopulations. This facilitates a fast and unbiased selection of training examples. For the generation of single cell images, the center of each cell was determined by its nuclear staining via the software CellProfiler (see above). Around each nuclei-center, a 50x50 pixel (or 39.5x39.5 µm) wide subimage was generated across all 5 measured channels. Single-cell sub-images were then manually annotated and sorted for their respective class using custom Matlab scripts. For training and validation of the discovery cohort CNN, a dataset of 89483 cells was manually annotated (containing both multiplexed and control staining cells). In the separate test datasets, each donor-associated set is independently split in multiplexed and control staining cells, resulting in a total of 30 independent test-datasets with each 100 cells per class. This test-setup allows inferring the network performance towards each donor, experiment and staining type independently.

Discovery cohort (10 donors): A 17-layer deep convolutional neural network with an adapted ‘Alex-Net’ architecture (Krizhevsky, Sutskever, and Hinton 2012) with 50x50 pixel and 5 channel input images was used. Before training, the labeled 8-class dataset was randomly split in a training set containing 90% and a validation set with the remaining 10% of all images. Network-layers weights and biases were initialized randomly before the CNN network was trained. Networks were trained up to 20 epochs with a mini batch size of 512 images. The learning rate was fixed to 0.0001. To avoid overfitting, L2 regularization with 0.005 was applied. Furthermore, in each iteration, input images were randomly rotated in 45-degree steps with an additional possibility to be also flipped vertically or horizontally. Performance of the trained networked was tested on the separate test-sets of staining control and multiplexed images of all 15 donors. Stochastic gradient descent with momentum of 0.9 is defined as the optimization algorithm. Finally, we trained 20 differently initialized networks with differently split training and validation sets. For the final classification of the complete unlabeled dataset the best performing network was used. As in the generation of the labeled dataset, 50x50 pixel sub-images around each nuclei-center were generated. Cells closer than 25 pixels to the border of an image were excluded from classification.

Validation cohort (15 donors): A 71-layer deep convolutional neural network with an adapted ResNet architecture (He et al. 2016) with 48x48 pixel and 5 channel input images was used. Before classification and training, all intensity values were first log_10_ transformed and then channel-wise normalized to a 0 to 1 range. The 8-class CNN was trained using randomly initialized weights and biases and the adaptive learning rate optimization ‘ADAM’. The network was trained for 20 epochs with an initial learning rate of 0.001 which was dropped every 5 epochs with a factor of 0.1. Furthermore, a mini batch size of 512 images and L2 regularization with 0.001 was applied. To further strengthen generalization, input images were augmented in each iteration. Here images were randomly rotated in 45-degree steps with an additional possibility to be also flipped vertically or horizontally. To block an over-reliance on absolute intensity values, channel intensity shifts were simulated via a multiplication with a random fixed factor. This used factor was randomly drawn out of a normal distribution with a mean of 1 and a standard deviation of 0.2. Furthermore, images were augmented with random noise (specifically salt and pepper noise, speckle noise, gaussian noise or image blurring). In all CNN classifications, 48x48 pixel sub-images around each nuclei-center were generated. Cells closer than 24 pixels to the border of an image were excluded from all classifications.

Label-free T-cell activation (T_ACT_) classier: Convolutional neural networks and single cell images were generated as described above. The labelled training and validation dataset comprised a total of 8862 cells (1:2 T_ACT_ :T_CON_ ratio). CNNs were trained with a mini batch size of 200 images to a maximum of 100 epochs, which could be terminated if validation loss was greater than the previous smallest loss for five consecutive times. Additionally, the images were randomly rotated by 45-degrees and mirrored vertically or horizontally per iteration to limit orientation bias towards polarised Tact cells. The CNN performance was assessed by classifying 1107 test cells (1:2 T_ACT_ Tact:T_CON_Tc ratio) that had neither been used in CNN training nor in validation.

#### RNA sequencing

T-cell isolation and RNA extraction: T-cells were isolated from fresh PMBCs directly after obtaining them via density centrifugation, as described above. Isolation was performed via a column based extraction method with CD3 Microbeads as described in the manufacturer’s instructions (Miltenyi Biotec). RNA extraction of the isolated cells was performed with a Quick-RNA MiniPrep Kit by Zymo according to manufacturer’s instructions.

RNA sequencing: RNA sequencing was performed by the Functional Genomics Center Zurich. In short, cDNA libraries were obtained according to protocols published by Picelli et al, 2014(Picelli et al. 2014). Illumina library was obtained via tagmentation using Illumina Nextera Kit. All samples were sequenced in a single run on a NovaSeq6000 (single read, 100bp, depth 20 Mio reads per sample).

Data processing and normalization: Illumina adapters, sequences of poor quality as well as polyA and polyT sequences were removed from the raw reads using TrimGalore v.0.6.0 with cutadapt v.2.0 prior to alignment. Reads were then aligned to the human reference genome GRCh38, v93 (Ensembl) using STAR v. 2.5.3a. Reads per gene were counted using the –quantMode GeneCounts flag in STAR. Gene counts below a threshold of 20 raw counts were filtered and raw counts were normalized (DESeq2(Love, Huber, and Anders 2014)). Only transcripts annotated as ‘protein coding’ or ‘long non-coding RNA’ were considered in the subsequent analysis.

### Statistical analysis

Significance calculation: If not stated otherwise all significance scores were calculated based on a two-tailed Student’s *t*-test with mean 0.

Cell-cell interaction analysis: For cell-cell interaction analysis, a simplified version of Vladimer et. al., 2017 (Vladimer et al. 2017) interaction method was used. Here, cell-cell interaction analysis was conducted over all different image sites within the same well. Cells were scored as interacting if their nuclear centroids were within a euclidean distance of 40 pixels. To calculate the interaction-score of a cell with type A interacting with a cell of type B, we first calculated specific interactions and total interactions per well. We define specific-interactions, as the total count of “B”-cells within the defined radius around a cell of type “A”. Total-interactions are considered as the total count of all interacting cells in that well. To calculate the final interaction score, specific-interactions were divided by the product of (the fraction of type A cells of all cells) × (the fraction of type B cells of all cells) × total-interactions. In contrast to the previously published method, this approach is simplified as the interactions scores are non-directed, which reduces the number of edges from 72 to 36. Mean interaction score over all replicates was calculated, log_2_-transformed and normalized towards its respective control (see Supplementary Table 2).

t-Distributed Stochastic Neighbor Embedding (t-SNE): All t-SNE visualizations were calculated on the –log_10_(class-probability matrices). In the t-SNE calculation a mahalanobis distance metric, a perplexity of 30, and an exaggeration parameter of 4 was applied. To reduce calculation time, the Barnes-Hut algorithm with a theta of 0.5 was used.

Local enrichment analysis (LEA): To calculate whether a certain condition displays local enrichment in the 8-dimensional class probability space, we developed local enrichment analysis by hypergeometric testing or rank-based correlation (LEA). Here, we probe the local neighborhood around each single cell, which is defined as the k-nearest neighbors in the original CNN class probability space. For discrete variables (such as donor identity), we calculate the probability to randomly find at least *n* cells of condition *X* in a certain neighborhood using a hypergeometric cumulative distribution function. This takes into account the total number of cells in the probed neighborhood, the total number of cells in the tested class probability space, and the total number of cells of condition *X*. In case of continuous variables (like donor age or gene transcript counts), the relative fraction of cells of each donor in the probed local neighborhood is calculated. The fractions are then correlated (Spearman’s rank correlation) with a continuous variable and the significance of the correlation is calculated. In both cases, the enrichment-probability is assigned to center-cell of the probed region and the approach is iterated for each single cell in the selected n-dimensional space. If not stated otherwise, neighborhoods were defined as k = 400 nearest neighbours for figures 2-3 and S2-S4 and k=200 for the T-cell figures 4-5 and S5-S6. P-values were corrected for multiple testing, i.e. by the number of total cells (i.e. tests) in the analysis.

Pathway enrichment analysis: Pathway annotations were obtained utilizing the David Database (Huang, Sherman, and Lempicki 2009). Gene enrichments per single cell were calculated via LEA (see above). To calculate pathway enrichments per single cell the LEA gene enrichments of all genes belonging to a certain pathway annotation were compared against the enrichment of all other genes. Significance scores were calculated based on a two-tailed Student’s *t*-test and directionality was calculated by the difference of the means of both populations.

### Data and Code availability

Further information and requests for resources and reagents should be directed to and will be fulfilled by the Lead Contact, Berend Snijder. This study did not create new unique reagents and all used reagents are commercially available.

CNN training and test datasets as well as the custom algorithm for local enrichment analysis by hypergeometric testing (LEA) will be available upon publication of this manuscript. The CNN dataset and relevant metadata is additionally available at the FAIR principles <u>(Wilkinson et al. 2016)</u> compliant repository https://doi.org/10.3929/ethz-b-000343106. Raw image data is available from the Lead Contact, upon request. T-cell RNA-seq measurements used in the study are available at https://www.ncbi.nlm.nih.gov/geo/query/acc.cgi?acc=GSE155093.

---

**Figure S1.**
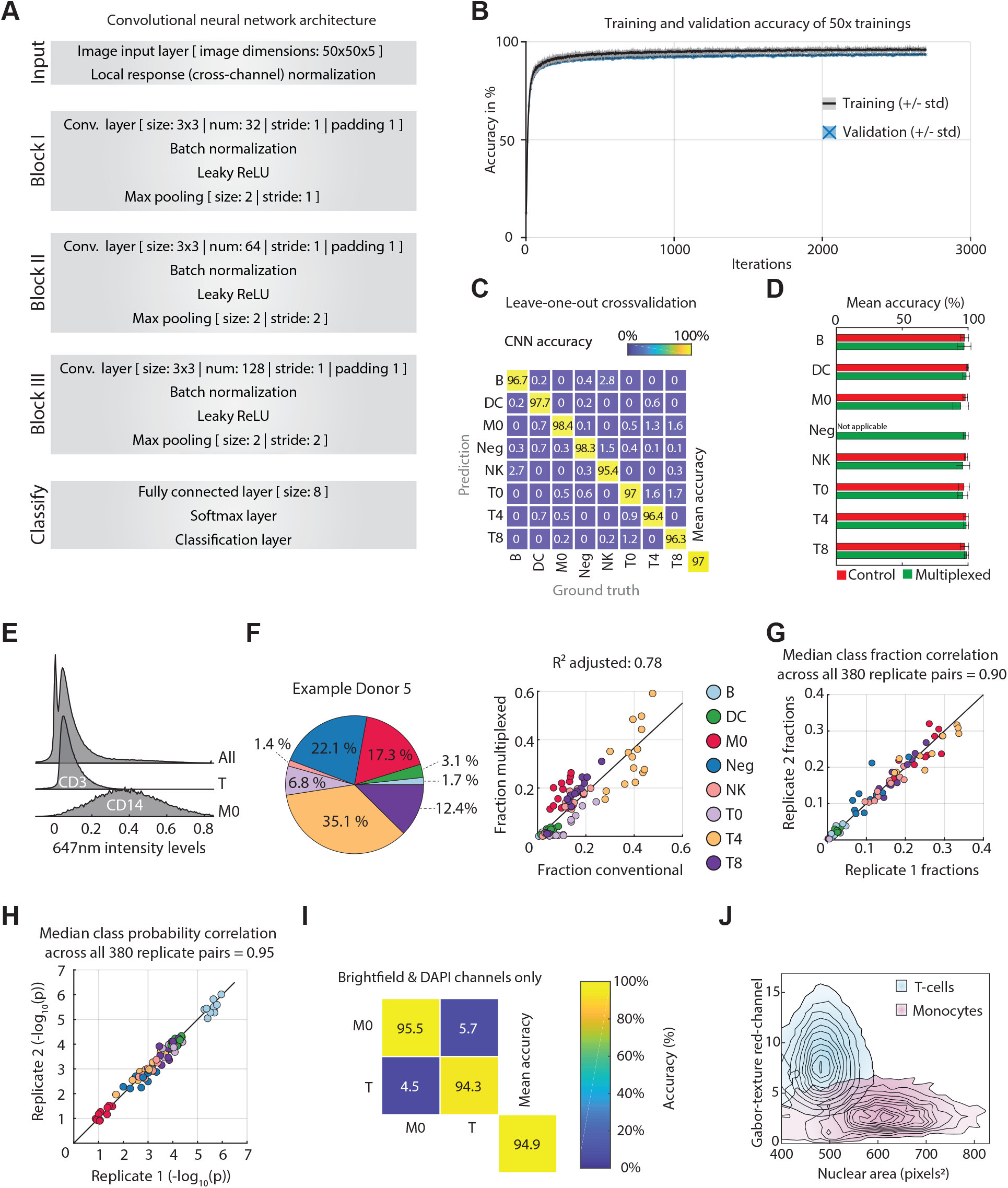
**(A)** Overview of the convolutional neural network architecture. **(B)** Average and standard deviation of training and validation accuracies over 20 randomly initialized CNN instances. Validation set represents 10% of the initial training set (n=8948). Network training after 20 epoches. **(C)** Confusion matrix of CNN performance on a leave-one-out cross validation per donor. The CNN was trained with 14 donors and subsequently tested on an unseen donor not included in the training dataset. The confusion matrix shows the mean accuracies after iterating across all donors. **(D)** Comparison of prediction accuracy on conventionally stained and multiplexed cells. Bar plots show the mean accuracy (in %) and the standard deviation of the CNN prediction across all donors individually per class and staining. **(E)** Distribution of 647nm intensity levels across all cells (upper), classified CD3^+^ T-cells (middle) and classified CD14^+^ monocytes (lowest) of Donor 1. A cell class probability threshold of 0.8 was applied. **(F)** Population percentages for Donor 5 (left) and class fraction comparison of conventionally stained and multiplexed cells across 15 healthy donors. Negative cell class is excluded due to its unavailability in conventional stainings. **(G)** Class fraction comparison of two single replicates (plate wells) across all 10 donors. Each dot corresponds to a replicate pair from a single donor. Color indicates the cell type. The median pairwise correlation across all technical replicates is indicated. **(H)** Median class probability comparison of two single replicates (plate wells) across all 15 donors. Shown statistic depicts the median class probability correlation of all pairwise replicate combinations per donor across two individual 384 well plates. **(I)** Confusion matrix of CNN performance on brightfield and DAPI channels only. An adapted CNN architecture (2-channel input and 2 class output) was trained with 1900 2-channel images of T-cells and monocytes. Network performance was evaluated in the curated test set containing 750 cells per class. **(J)** Comparison of selected morphological and staining-pattern parameters divergent between T-cells and monocytes. Conventionally stained T-cells and monocytes from Donor 1 were identified by immunofluorescence gating for CD3 and CD14, respectively. Morphological features were extracted by CellProfiler.

**Figure S2.**
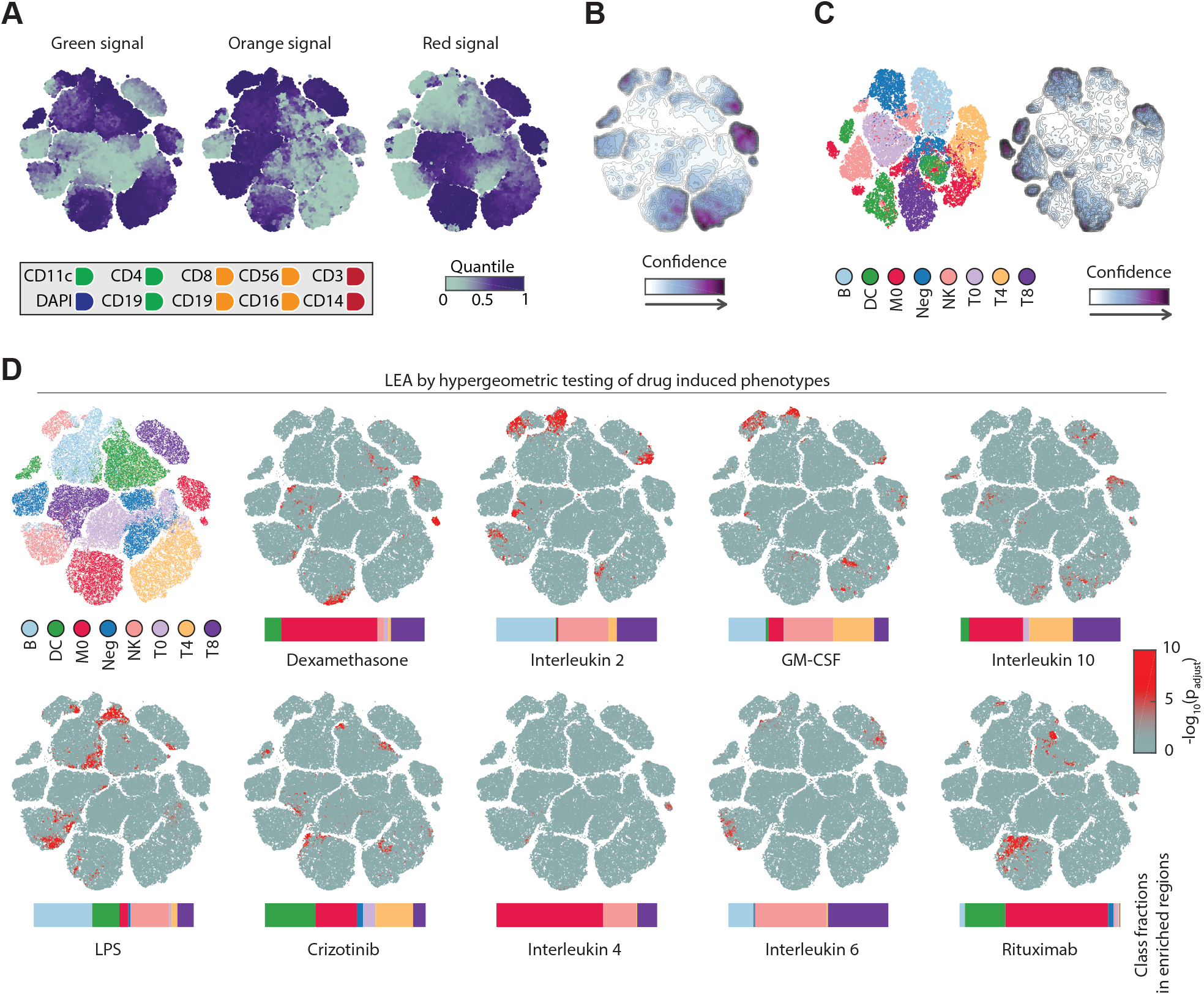
**(A)** Selected single-cell features mapped onto the same t-SNE map as depicted in Figure 2D. Median value of overlapping data points is calculated and color is assigned accordingly. Points are plotted in order of intensity, from highest to lowest. **(B)** Associated CNN probability contour plot of the phenotypic landscape of the immune system depicted in Figure 2D. **(C)** Left: Phenotypic landscape of the immune system across ten healthy donors. t-SNE embedding of the 8-class CNN probabilities without a confidence threshold of up to 1000 randomly subsampled multiplexed cells per class and per donor. Right: Associated CNN probability contour plot of the phenotypic landscape depicted left. **(D)** LEAs visualized by t-SNE of drug induced phenotypes. Horizontal bar graphs indicate the class fractions in enriched regions (at p_adjust_ < 0.01).

**Figure S3.**
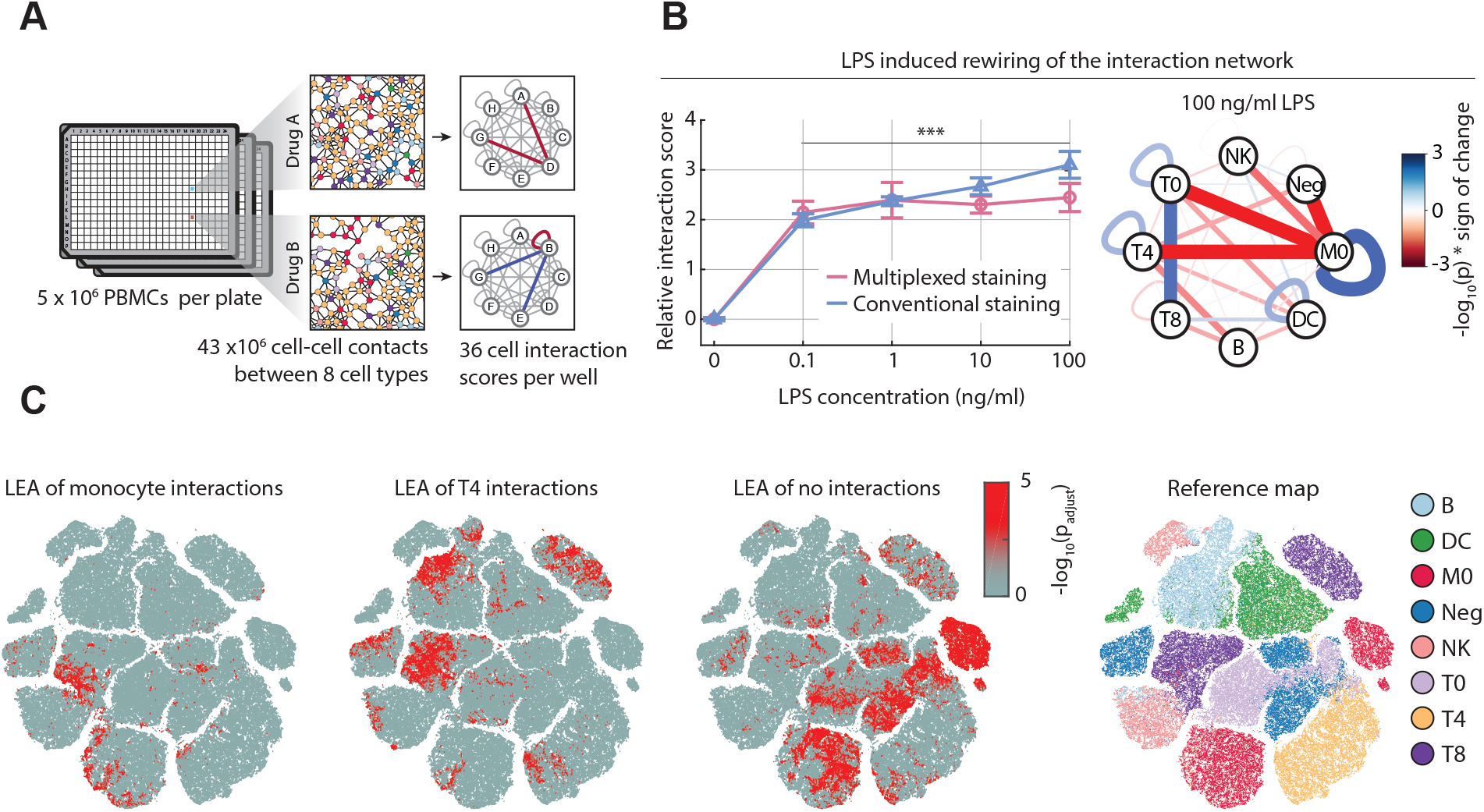
**(A)** Overview of cell-cell contact analysis over five million PBMCs. Cell-to-cell interaction networks between eight different immune cell populations with a total of 36 cell type interactions were generated per well, and compared across treatments. **(B)** LPS-induced rewiring of the cell-to-cell interaction network. Relative monocyte-to-monocyte interaction scores of multiplexed and conventionally stained wells as a function of increasing LPS concentration (left). Mean interaction score across all replicates is calculated and normalized against control treatment. Example LPS interaction network for 100ng/ml LPS (right). Significance of interaction (−log_10_(p), multiplied times the sign of the phenotype (either positive or negative interaction score)). **(C)** LEA of cells with monocytes (left), T-cells (middle) or no-nearest neighbor (right).

**Figure S4.**
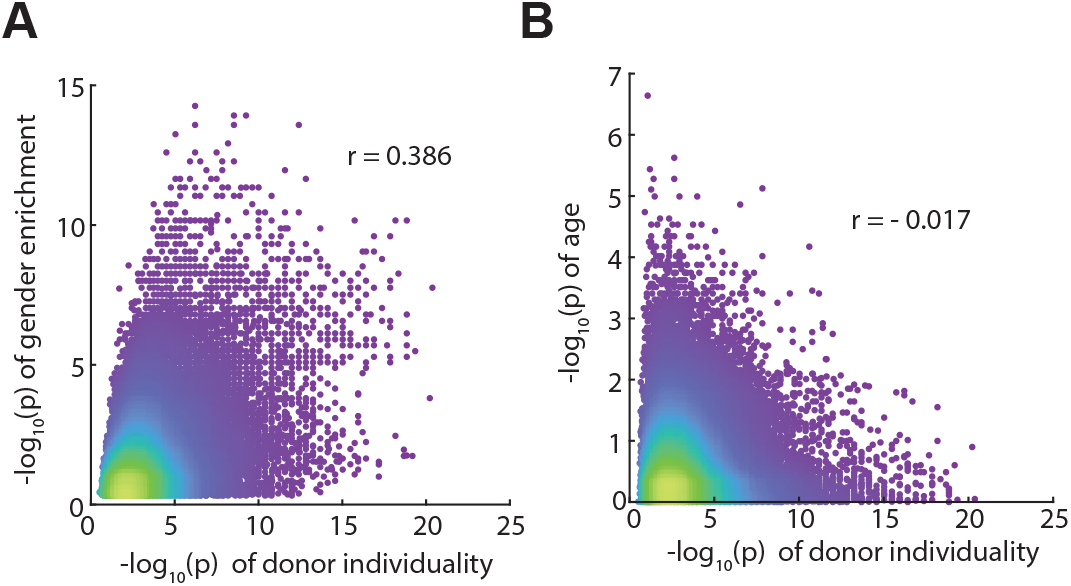
**(A)** Comparison of donor LEA enrichments vs gender LEA enrichments per single cell (as in Figure 3D). r values represent Pearson correlations. **(B)** Comparison of donor LEA enrichments vs age LEA enrichments per single cell (as in Figure 3D). r values represent Pearson correlations.

**Figure S5.**
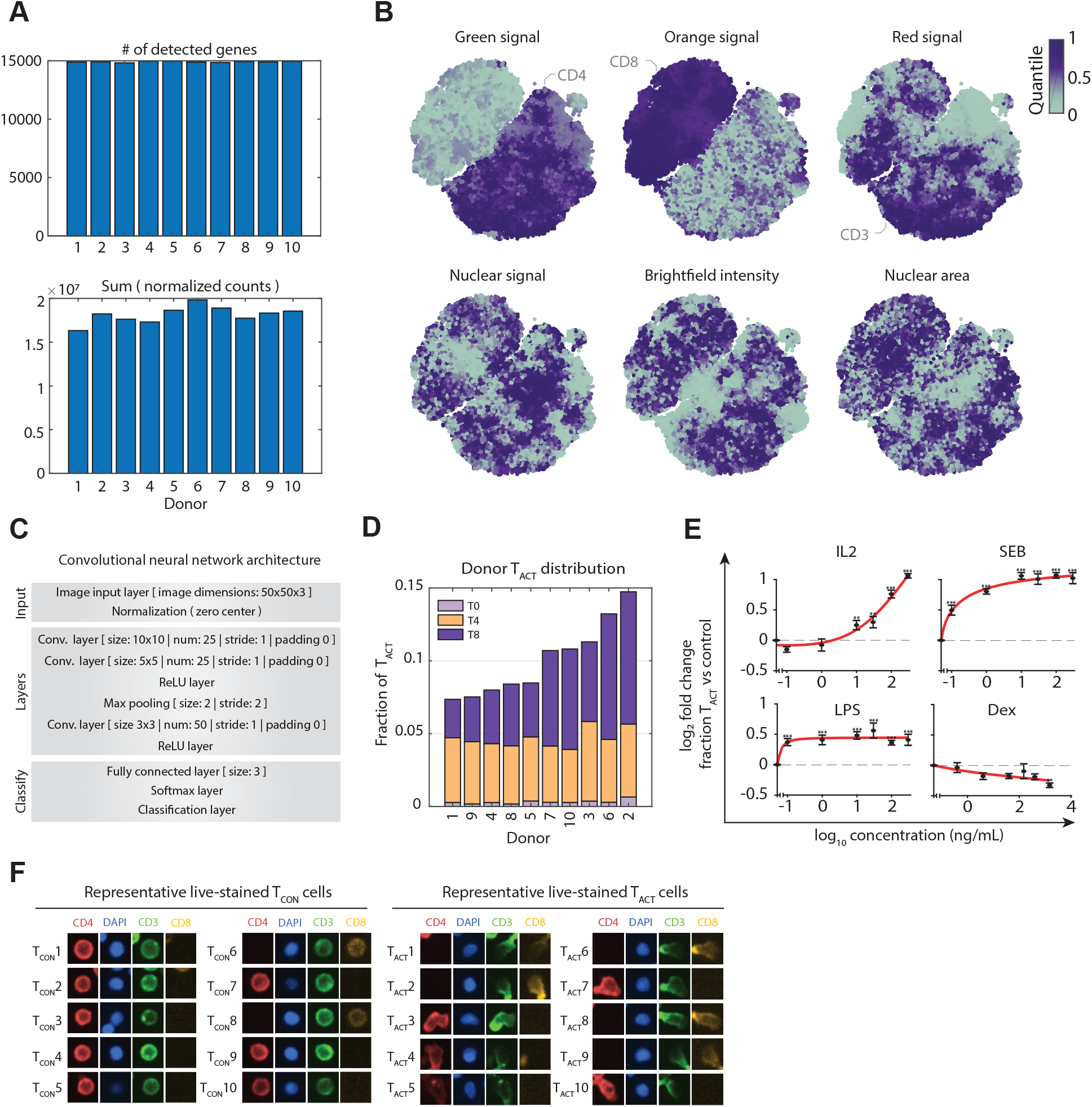
**(A)** Upper: Bar graphs indicate the number of detected transcripts (protein coding and long non-coding RNAs) after applying a threshold of 20 raw counts. Lower: Bar graphs indicated the sum of transcript counts after DESeq2 normalization (Love, Huber, and Anders 2014). **(B)** Selected single-cell features projected onto the t-SNE depicted in Figure 4A. Median value of overlapping data points is calculated and color is assigned accordingly. Points are plotted in ascending order with the lowest intensity on top. **(C)** Overview of the label-free T-cell activation (T_ACT_) convolutional neural network architecture. **(D)** Fraction of T_ACT_ cells per class and per donor. Stacked bar plots show the mean fraction of all T-cells per donor classified as T_ACT_, within their respective T-cell subclass (T0, T4 or T8) in control (DMSO) conditions. **(E)** Induction and suppression of the T_ACT_ cell phenotype by immunomodulatory agents. Plotted are the log_2_ fold changes of the mean fraction of T-cells classified as T_ACT_ across all wells of each drug condition compared to control treatments. Cells were incubated with immunomodulatory agents at 0.1, 1, 10, 30, 100 and 300 ng/ml. Error bars show the standard error of the mean across wells for each drug condition. A custom Hill function (adjusted to different minima and maxima) was used to fit the data (red line). **(E)** Representative live-stained T_ACT_ and T_CON_ cell morphologies. Crop-size is 15 x 15 µm.

**Figure S6.**
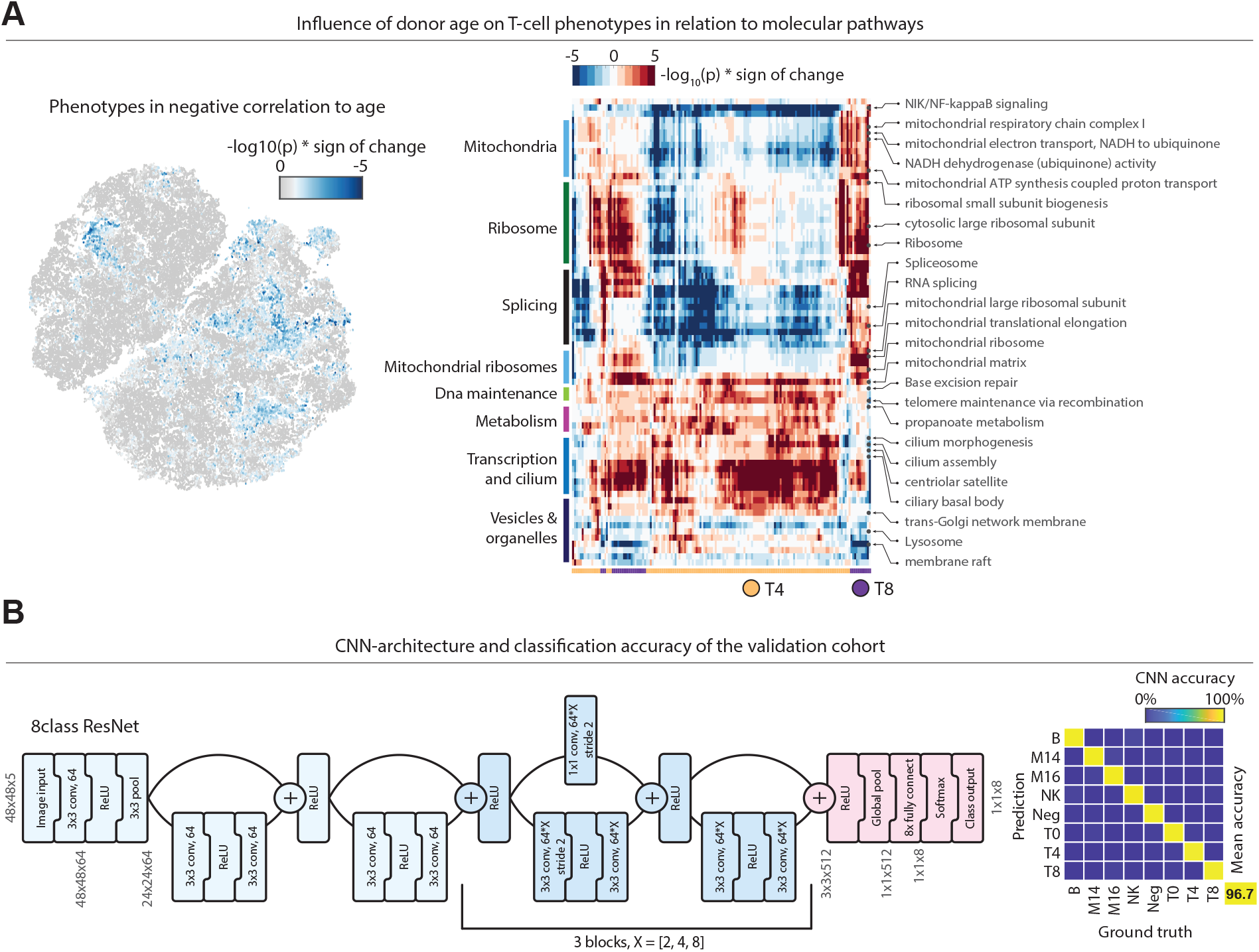
**(A)** Left: negative LEA associations with donor age projected onto the t-SNE (colored by - log_10_(p)). Right: Heatmap overview of all significantly enriched pathways in positive age-associated T-cells (-log_10_(p)<-5). Rows are annotations, columns are significantly age-associated cells. **(B)** Left: Schematic of the 8-class ResNet architecture used for the 15 donor validation cohort. Right: Confusion matrix of the CNN. CNN was tested on 1000 cells per class that the CNN did not see before.

**Supplementary Table 1.**

| Epitope | Vendour | Fluorophore | Clone | Host | Staining Control | Used in multiplexed | Lot |
| --- | --- | --- | --- | --- | --- | --- | --- |
| CD3 | BioLegend | Alexa Fluor 647 | UCHT1 | Mouse | 1 | yes | B246715 |
| CD4 | BioLegend | FITC | SK3 | Mouse | 1 | yes | B244280 |
| CD8 | BioLegend | Alexa Fluor 594 | RPA-T8 | Mouse | 1 | yes | B200099 |
| CD19 | BioLegend | FITC | SJ25C1 | Mouse | 3 | yes | B239447 |
| CD19 | BioLegend | PE | SJ25C1 | Mouse | 3 | yes | B237928 |
| CD56 | Beckman Coulter | PE | N901 | Mouse | 2 | yes | 52 |
| CD16 | BioLegend | PE | 3G8 | Mouse | 2 | yes | B238510 |
| CD14 | BioLegend | Alexa Fluor 647 | HCD14 | Mouse | 2&3 | yes | B260484 |
| CD11c | BioLegend | Alexa Fluor 488 | 3.9 | Mouse | 2&3 | yes | B209841 |
| CD3 | BioLegend | Alexa Fluor 488 | UCHT1 | Mouse | / | no | B278994 |
| CD4 | Biolegend | Alexa Fluor 647 | SK3 | Mouse | / | no | B293054 |
| CD8 | Biolegend | Alexa Fluor 594 | RPA-T8 | Mouse | / | no | B200099 |
| pNFkB p65 (Ser529) | eBioscience | PE | B33B4WP | Mouse | / | no | 4303324 |
| pERK1/2 (Thr202 Tyr204) | Thermo Fisher Scientific | PE | MILAN8R | Mouse | / | no | 4337535 |

**Supplementary Table 2.**

| Compound name | Vendour | Assay conc. | Carrier solution and control |
| --- | --- | --- | --- |
| Crizotinib | Sigma-Aldrich | 10uM | 1% DMSO |
|  |  | 1uM | 0.1% DMSO |
|  |  | 0.1uM | 0.01% DMSO |
|  |  | 0.01uM | 0.001% DMSO |
| Dexamethasone | Sigma-Aldrich | 400ng/ml | 1% DMSO |
|  |  | 40ng/ml | 0.1% DMSO |
|  |  | 4ng/ml | 0.01% DMSO |
|  |  | 0.4ng/ml | 0.001% DMSO |
| Lipopolysaccharide from Escherichia coli | Sigma-Aldrich | 100ng/ml | PBS |
|  |  | 10ng/ml | PBS |
|  |  | 1ng/ml | PBS |
|  |  | 0.1ng/ml | PBS |
| Rituximab | Absolute antibody | 1ug/ml | PBS |
|  |  | 0.5ug/ml | PBS |
|  |  | 0.1ug/ml | PBS |
|  |  | 0.05ug/ml | PBS |
| Recombinant Human IL-2 | PeproTech | 100ng/ml | 0.001% (w/v) BSA in PBS |
|  |  | 10ng/ml | 0.0001% (w/v) BSA in PBS |
|  |  | 1ng/ml | 0.00001% (w/v) BSA in PBS |
|  |  | 0.1ng/ml | 0.000001% (w/v) BSA in PBS |
| Recombinant Human IL-4 | PeproTech | 100ng/ml | 0.001% (w/v) BSA in PBS |
|  |  | 10ng/ml | 0.0001% (w/v) BSA in PBS |
|  |  | 1ng/ml | 0.00001% (w/v) BSA in PBS |
|  |  | 0.1ng/ml | 0.000001% (w/v) BSA in PBS |
| Recombinant Human IL-6 | PeproTech | 100ng/ml | 0.001% (w/v) BSA in PBS |
|  |  | 10ng/ml | 0.0001% (w/v) BSA in PBS |
|  |  | 1ng/ml | 0.00001% (w/v) BSA in PBS |
|  |  | 0.1ng/ml | 0.000001% (w/v) BSA in PBS |
| Recombinant Human IL-10 | PeproTech | 100ng/ml | 0.001% (w/v) BSA in PBS |
|  |  | 10ng/ml | 0.0001% (w/v) BSA in PBS |
|  |  | 1ng/ml | 0.00001% (w/v) BSA in PBS |
|  |  | 0.1ng/ml | 0.000001% (w/v) BSA in PBS |
| Recombinant Human G-MCSF | PeproTech | 100ng/ml | 0.001% (w/v) BSA in PBS |
|  |  | 10ng/ml | 0.0001% (w/v) BSA in PBS |
|  |  | 1ng/ml | 0.00001% (w/v) BSA in PBS |
|  |  | 0.1ng/ml | 0.000001% (w/v) BSA in PBS |

**Supplementary Table 3.**

| Discovery cohort (Figure 1, Figure 2 A-B, Figure 3, Figure 4 A-G, Figure 5 A-B) |  |  |  |  |  |  |  |
| --- | --- | --- | --- | --- | --- | --- | --- |
| Whole Blood Donor | Year of birth | Blood type | weight in kg | Height in cm | Gender | Blood pressure | Hb level |
| 1 | 1999 | A+ | 68 | 174 | m | 128/70 | 156 |
| 2 | 1987 | A+ | 66 | 170 | m | 173/99 | 147 |
| 3 | 1990 | A+ | 50 | 163 | f | 114/80 | 148 |
| 4 | 2000 | A+ | 63 | 168 | f | 122/74 | 133 |
| 5 | 1968 | O- | 58 | 165 | f | 158/90 | 151 |
| 6 | 1974 | B+ | 95 | 176 | m | 156/98 | 177 |
| 7 | 1950 | O+ | 72 | 175 | f | 126/78 | 130 |
| 8 | 1967 | A+ | 95 | 178 | m | 146/86 | 161 |
| 9 | 1967 | O+ | 80 | 180 | m | 160/106 | 167 |
| 10 | 1965 | A+ | 80 | 180 | m | 130/82 | 167 |

| Immuno-modulatory screen (Figure 2 C-F) |  |  |  |  |  |  |  |
| --- | --- | --- | --- | --- | --- | --- | --- |
| Buffy coat donor | Year of birth | Blood type | weight in kg | Height in cm | Gender | Blood pressure | Hb level |
| 1 | 1968 | AB+ | 73 | 180 | m | 146/92 | 153 |

| Validation cohort (Figure 4H, Figure 5C) |  |  |  |  |  |  |  |
| --- | --- | --- | --- | --- | --- | --- | --- |
| Buffy coat Donor | Year of birth | Blood type | weight in kg | Height in cm | Gender | Blood pressure | Hb level |
| 1 | 1994 | B+ | 65 | 183 | m | 135/77 | 152 |
| 2 | 1970 | B+ | 69 | 177 | f | 148/92 | 147 |
| 3 | 1996 | B+ | 75 | 192 | m | 138/91 | 157 |
| 4 | 1994 | B+ | 85 | 180 | m | 124/74 | 159 |
| 5 | 1953 | B- | 65 | 178 | m | 142/95 | 168 |
| 6 | 1971 | AB- | 65 | 171 | f | 114/72 | 141 |
| 7 | 1991 | B+ | 67 | 180 | f | 135/94 | 145 |
| 8 | 1963 | B+ | 65 | 157 | f | 146/99 | 157 |
| 9 | 1963 | B+ | \ | \ | m | 170/90 | 169 |
| 10 | 1953 | O- | 76 | 158 | f | 142/100 | 149 |
| 11 | 1999 | AB+ | 61 | 169 | f | 131/74 | 136 |
| 12 | 1960 | B+ | 90 | 182 | m | 155/98 | 151 |
| 13 | 1999 | B+ | 66 | 173 | m | 171/103 | 157 |
| 14 | 1967 | B+ | 80 | 182 | m | 162/102 | 174 |
| 15 | 1989 | AB- | 72 | 178 | f | 108/63 | 138 |

